# Population structure and diversity of common bean (*Phaseolus vulgaris* L.) landraces in the Peruvian Amazon

**DOI:** 10.1101/2025.09.04.674357

**Authors:** Jorge Tobaru, Marvin Barrera-Lozano, Carlos D. Vecco-Giove, Jorge Peláez, Mack Pinchi, Yesenia Vargas, Jorge Chuquillanqui, Julian Soto-Torres, Ronald Robles, David Saravia, Cesar Petroli, Jorge D. Etchevers, Rodomiro Ortiz, Raul Blas

## Abstract

Common bean (*Phaseolus vulgaris* L.) is a globally important legume with significant nutritional and agronomic value. Genetic diversity in Amazonian landraces remains undercharacterized, limiting their utilization in breeding programs. Here we analyzed 647 accessions from the Peruvian Amazon using 18 morphological traits and 23,050 high-quality DArTseq single nucleotide polymorphisms (SNPs). Our results revealed two major genetic groups corresponding to Andean and Mesoamerican gene pools, each subdivided into distinct subgroups, exhibiting high allelic richness and structured diversity. Despite abundant polymorphic loci, observed heterozygosity was low, consistent with the species’ self-pollinating nature. These findings highlight the Peruvian Amazon as a reservoir of unique genetic variation, underscoring the need for integrated in situ and ex situ conservation strategies. This germplasm offers valuable resources for breeding programs targeting tropical adaptation and resilience, contributing to sustainable agriculture and food security.

## Introduction

The common bean (*Phaseolus vulgaris* L.) an annual leguminous self-pollinated crop is the most widely cultivated species of the genus Phaseolus and a major source of dietary vegetal protein and income worldwide; which is a diploid (2n = 2x = 22) species and its genome size according to the first telomere-to-telomere (T2T) genome assembly was 560.30 Mb and predicted 29,925 protein-coding genes (Wang et al., 2025) [1]. The origin of *P. vulgaris* remains controversial, separating it from Mesoamerica and the Andean region. However, recent research using molecular markers, including SNPs, shows that this species’ origin is Mesoamerica [2] [3] [4] [5]. Genetic and phylogeographic evidence indicates three distinct wild gene pools: 1) Mesoamerica (Mexico, Central America, Colombia and Venezuela), 2) the Andes (southern Peru, Bolivia and Argentina), and 3) northern Peru/Ecuador; nevertheless, the three common bean genetic groups belong to the primary gene pool [6]. From those, two main domesticated groups emerged belonging to the Mesoamerican and Andean gene pools, this domestication process has been done independently in the Andes and Mesoamerica, which are associated with two Andean and Mesoamerican gen pools, respectively [7] [8]. Both types of beans were introduced to other continents such as Europe, Africa, and Asia, where their widespread acceptance has led to debate about whether Europe can be a secondary center of diversification [9].

At the present, it is a staple crop in many parts of the world, most widely cultivated legume species worldwide, mainly due to its nutritional composition, being rich in amino acids, minerals, and vitamins—especially lysine, iron, zinc and folic acid [10]; presents medicinal attributes such as the prevention of cardiovascular diseases [11]; and has an ecological importance, it fixes atmospheric nitrogen reducing all the environmental costs involved in the acquisition of chemical fertilizers [12]. For this reason, the United Nations declared in 2016 as “the international year of legumes” in its 68^th^ session [13], the common bean being within the legumes, considered one of the essential crops to promote food security. Common bean has thus become an important source of income and nutrients for small farmers in the Caribbean, Latin America, Asia, and Africa, where account for 77% of global production [14] [15] [16]. In the case of Brazil, a predominantly Amazonian region, the Mesoamerican gene pool is found in a greater proportion (79%) and the Andean gene pool in a lesser proportion (21%) [4].

In Peru, the common bean is cultivated across diverse agroecological zones, ranging from coastal valleys to highland and Amazonian regions. According to national statistics [17], production covers approximately 42,575 hectares annually, with San Martín, Piura, and Cajamarca being leading regions; with around 14% of this area located in the Amazon region—2.0% less than the area recorded in 2023. Beyond its economic relevance, the crop plays a crucial role in household nutrition and local food security. Despite its importance, most studies on common bean in Peru have focused on commercial varieties grown in the highlands and coastal regions, emphasizing agronomic practices and genetic improvement using both local and introduced varieties. Conversely, there is limited research conducted on the varieties cultivated by farmers in indigenous communities of the Peruvian Amazon rainforest. According to the Genesys database, from Peru holds 3,497 accessions of common beans, of which 2,838 (81%) are classified as traditional varieties, 516 (15%) are results from breeding programs, and 143 (4%) correspond to wild or near-wild forms [18]. The International Center for Tropical Agriculture (CIAT) holds the global reference collection, while the Peruvian National Institute for Agrarian Innovation (INIA) maintains the national bean germplasm bank, which officially has 58 accessions of beans and 146 accessions of popping bean mainly from the Andean region [19] [20]. Despite these efforts, landraces and wild relatives from the Peruvian Amazon remain underrepresented, leaving a gap in the documentation of local agrobiodiversity.

Research on cultivated common bean diversity in Peru remains scarce and typically focuses on specific types such as popping beans (“ñuña”, in the Quechua language) using phenotypic and molecular markers [21] [22] or on specific localities [23]. [24] studied the cultivated and sociocultural diversity of beans in the Amazon region of central Peru, documenting practices across several Indigenous communities (ethnic groups). More recently, [25] conducted an agromorphological characterization of 58 Phaseolus spp. accessions in the Amazonas region, identifying promising genotypes with yields exceeding 2.77 t/ha with potential use in breeding programs. Genotyping using DArTseq SNP has been used in other legume crops as cowpea [26] and chickpea [27] and common bean of Ethiopian germplasm [28]. However, such studies remain largely unexplored in common bean from Peru. The Peruvian Amazon harbors a wide array of common bean landraces and wild relatives, which are essential for preserving genetic diversity. The adaptation of common bean to the tropical rainforest conditions is particularly important due to the challenges associated with high humidity, variable soil fertility, and significant pathogen pressure. However, no systematic or representative collection has been established from the Peruvian Amazon, nor has a comprehensive analysis of the genetic diversity across its rainforest regions been conducted. As a result, little is known about the genetic diversity and origins of these varieties, which may represent an untapped reservoir of alleles for crop improvement. This study aimed to assess the genetic diversity of common bean landraces collected in the Peruvian Amazon using both morphological descriptors and genome-wide SNP markers (DArTseq). By combining phenotypic and molecular approaches, we sought to clarify the structure of local germplasm and to generate information that supports both conservation strategies and future breeding efforts.

## Materials and Methods

### Sample Collection

A total of 1,025 common bean (*P. vulgaris*) accessions were recently collected from 10 rainforest regions of Peru: Amazonas, Cajamarca, San Martín, Huánuco, La Libertad, Loreto, Ucayali, Cerro de Pasco, Junín, and Cusco across the Peruvian Amazon, including Indigenous farming systems and traditional local markets (Figure 1). From this collection, a subset of 633 accessions— representative from Amazonian region—was selected for morphological and molecular characterization. Also, it was added 34 accessions for genotyping as a reference genetic material belonging to both Mesoamerican and Andean gene pools, looking for the relationships between accessions and to help to define gene pool group; from them 26 are from Peru (12 from Coast, 14 from Highlands), 7 from CIAT (2 Andean y 5 Mesoamerican gene pool), and 1 from Mexico which was considered as a reference for Mesoamerican gene pool.

**Figure 1.**
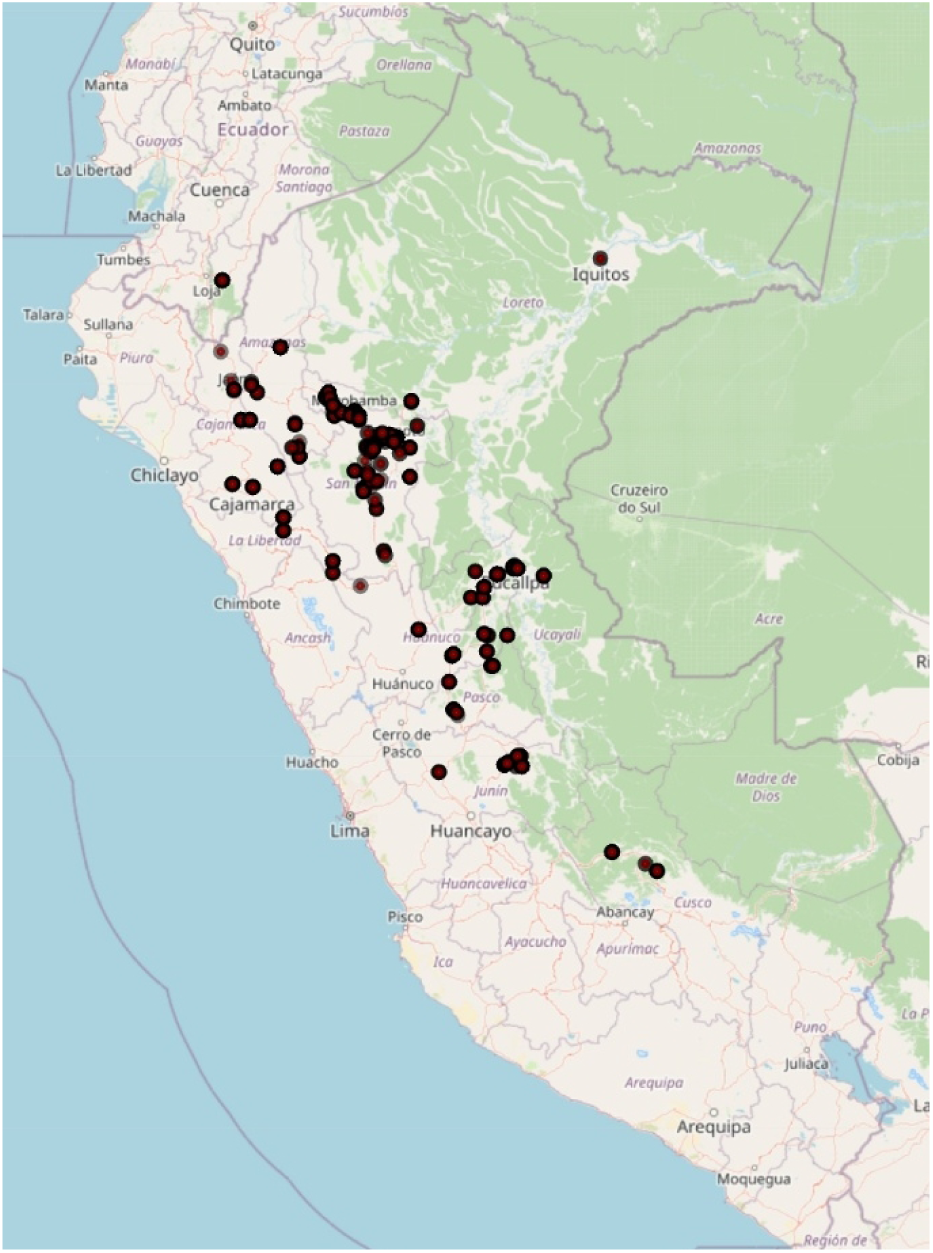
Distribution of Common bean accession from Peruvian Amazonia.

### Morphological Characterization

The experimentation was conducted in 2024 cropping seasons at the experimental field of Universidad Nacional Agraria La Molina (UNALM), Lima, Peru. The density of sowing was in a 2.5 m row: 35 cm between plants and 50 cm between row, with up to three seeds per hole (in total by experimental unit, 7 holes and 21 plants). Data collection followed the *Phaseolus vulgaris* descriptors established by the International Board for Plant Genetic Resources [29]. Traits evaluated included plant growth habit, and pod characteristics (shape, color, size, and presence of sutures or fiber). Throughout the growing cycle, a total of 18 traits were recorded—3 quantitative and 15 qualitative—covering vegetative, floral, pod, and seed characteristics (Table 1); such as seed shape, size, color, and plant growth habit were evaluated according to standard descriptors. For each accession, five plants were evaluated, and from each plant, five pods and ten seeds were analyzed. Pods were collected at full physiological maturity.

**Table 1.** Quantitative and qualitative traits recorded in the *Phaseolus vulgaris* accessions. IBPGR— International Board for Plant Genetic Resources.

| Order | Descriptor title | Descriptor data type | Codes titles |
| --- | --- | --- | --- |
| 1 | Plant type/<br>Growth habit | Coded | 1 Determinate bush; 2 Indeterminate bush; 3 Indeterminate semi-climber or prostrate (with many lateral guides); 4 Indeterminate climber |
| 2 | Days to flowering | Numeric | Number of days from emergence to stage where 50% of plants have set flowers |
| 3 | Color of standard | Coded | In freshly opened flowers: 1 White; 2 Green; 3 Lilac; 4 White with lilac edge; 5 White with red stripes; 6 Dark lilac with purple outer; 7 Dark lilac with purplish spots; 8 Carmine red; 9 Purple; 10 Light lilac; 11 Dark lilac |
| 4 | Color of wings | Coded | In freshly opened flowers: 1 White; 2 Green; 3 Lilac; 4 White with carmine stripes; 5 Strongly veined in red to dark lilac; 6 Plain red to dark lilac; 7 Lilac with dark lilac veins; 8 Purple; 9 Light lilac; 10 Dark lilac |
| 5 | Pod color | Coded | From fully expanded immature pod: 1 Dark purple; 2 Carmine red; 3 Purple stripe on green; 4 Carmine stripe on green; 5 Pale red stripe on green; 6 Dark pink; 7 Normal green; 8 Shiny green; 9 Dull green to silver grey; 10 Golden or deep yellow; 11 Pale yellow to white. |
| 6 | Pod length | Numeric | Average length in centimetres of the largest fully expanded immature pods from 10 random normal plants |
| 7 | Pod cross-section | Coded | From fully expanded immature pod: 1 Very flat, with probable; 2 Pear shaped; 3 Round elliptic; 4 Figure of eight |
| 8 | Pod curvature | Scale | Fully expanded immature pod: 3 Straight; 5 Slightly curved; 7 Curved; 9 Recurving |
| 9 | Locules per pod | Numeric | Number of locules from longest pod of 10 random normal plants |
| 10 | Pod wall fibers | Scale | 3 Strongly contracting (at dry maturity adhering around seed)<br>5 Leathery podded (dry pods will not spontaneously open)<br>7 Excessive shattering (with strong twisting of dry pods) |
| 11 | Pod apex position | Coded | 1 = marginal; 2 = non-marginal; 3 = other. |
| 12 | Pod apex orientation | Coded | 1 = upwards (dorsal direction); 2 = straight; 3 = downwards (ventral direction). |
| 13 | Pod color at the physiological maturity | Coded | 1 Dark purple; 9 Silver; 10 Golden or deep yellow; 11 Light yellow to white; 12 Deep yellow with purple mottling or strips; 13 Pale yellow with purple mottling or stripes; 14 Pale yellow with red mottling or stripes |
| 14 | Seed coat pattern | Coded | 1 Constant mottled; 2 Striped; 3 Rhomboid spotted; 4 Speckled; 5 Circular mottling; 6 Marginal color pattern; 7 Broad striped; 8 Bicolor; 9 Spotted bicolor; 10 Absent or whole color |
| 15 | Seed coat lighter color | Text | 1 Black; 2 Brown, pale to dark; 3 Maroon; 4 Grey, brownish to greenish; 5 Yellow to greenish; 6 Pale-cream to buff; 7 Pure white; 8 Whitish; 12 Red; 13 Pink; 14 Purple; 15 Carmine red or wine; 16 Dark yellow |
| 16 | Seed coat darker color | Coded | 1 Black; 2 Brown, pale to dark; 3 Maroon; 4 Grey, brownish to greenish; 5 Yellow to greenish; 6 Pale-cream to buff; 7 Pure white; 8 Whitish; 12 Red; 13 Pink; 14 Purple; 15 Carmine red or wine; 16 Dark yellow |
| 17 | Seed brightness | Scale | 3 Matt; 5 Medium; 7 Shiny |
| 18 | Seed shape | Coded | Taken from middle of pod :1 Round; 2 Oval; 3 Cuboid; 4 Kidney shaped; 5 Truncate fastigate |

### Morphological data analysis

Quantitative traits were summarized as boxplot, mean, and minimum and maximum values. The qualitative characteristics were expressed on scales, and then graphically represented in the shape of Histogram charts. A principal component analysis (PCA) was used to examine the association between the analyzed traits and the similarity among accessions. PCA was performed with all quantitative traits and qualitative categorized values. Additionally, for the quantitative traits, a Euclidean distance matrix based on standardized data was computed for clustering analysis by using the UPGMA (unweighted pair group method with arithmetic mean) method. Those analysis was made using R version 4.5.1 [30] and RStudio version 2025.05.1+513 [31], programming language for statistical computing and data visualization, using R Base and packages as “FactoMineR” [32], “factoextra” [33], “ggplot2” [34], and “randomForest” [35].

### Genotyping

#### DNA isolation and sequencing

Seeds of each common bean accessions were germinated in a shade house. Leaf samples were collected from two-week-old seedlings, immediately frozen at −80 °C, and used for DNA extraction. Genomic DNA was extracted from frozen leaf tissue using CTAB protocol. DNA quality and concentration were initially assessed using a NanoDrop spectrophotometer and subsequently quantified with a Qubit fluorometer (Life Technologies, Carlsbad, CA, USA). DNA Integrity was verified via 0.8% agarose gel electrophoresis. The DNA concentration was adjusted to 50–100 ηg µl−1 and samples were submitted for genotyping using high-throughput genotyping DArTSeq™ technology platform at SAGA-CIMMYT(Mexico), following the protocol described [36] and [37]. This technology uses a complexity reduction approach based on a digestion/ligation reaction, employing a combination of two restriction enzymes, PstI and MsI, to enrich genomic representations with single-copy sequences. PstI-compatible adapters were ligated for subsequent PCR amplification, along with sample-specific barcodes added to the digested fragments. Equimolar amounts of amplified products from each sample were pooled and sequenced on an Illumina Novaseq 6000 platform. Sequence data were processed using proprietary DArT analytical pipelines to filter low-quality reads and assign sequences to individual samples based on barcode information.

#### SNP calling and genotypic data filtering

Molecular markers discovery was performed using an analytical pipeline (DArTsoft14) developed by DArT company, this software identifies two types of markers: SilicoDArT (presence/absence) and SNP markers [38]. Marker scoring and reproducibility were assessed using technical replicates, and only markers with high call rates and reproducibility were retained. Both markers were aligned to the Phaseolus vulgaris YP4 reference genome [1] to identify the chromosome positions. Initially, we received 80,079 DArTSeq-derived SNP markers from SAGA-CIMMYT, which were polymorphic across common bean genotypes analyzed. Markers with unknown position were first removed from the analysis. Marker parameters such as call rate and minimum allele frequency (MAF) were calculated using dartR package in R version 4.5.1 [39]. Accordingly, SNPs markers with ≥ 50% call rate and MAF of ≥ 1% were retained for further analysis. In this study, genotypes with missing data above ≥ 30% were removed from the analysis. For SilicoDArT markers (presence / absence), the initial report of 462,250 markers was filtered to retain markers with call rate ≥ 90% and MAF of ≥ 1%. Markers with unknown position were also removed from the final analysis.

#### Genetic relationships and diversity metrics

Pairwise genetic distances among accessions were calculated using the scaled Euclidean distance method, as implemented in the gl.dist.ind function of the dartR package. Genetic diversity indices, including expected heterozygosity (He), observed heterozygosity, polymorphic information content (PIC), major allele frequency, and inbreeding coefficient (Fis), were computed using functions also available in dartR package.

To infer relationships among accessions, we constructed a neighbor-joining tree, based on the previous euclidean distance matrix, in DARwin v6.2.1. [40]. Principal Coordinate Analysis (PCoA) was conducted using the gl.pcoa function in dartR to visualize genetic relationships among accessions in reduced dimensions.

#### Population structure and genetic diversity analysis

Population structure was inferred using fastSTRUCTURE, a framework for inferring population structure from a large SNP genotype dataset [41]. Analyses were conducted for a range of K values (number of assumed populations) from 1 to 10. The ancestry proportions for each individual (Q-matrix) were visualized alongside the neighbor-joining tree and combined with the geographical passport data to contextualize cluster assignments.

## Results

### Nominal variability of common bean

Based on passport data collected during fieldwork a total of 73 different common names were recorded for common bean from Peruvian Amazonia. The ten most frequently cited varieties were Panamito, Waska, Ashpa, Pinto, Canario, Poroto, Ñuña (popping bean), Shingo, Manteca and Awisho (Table 2). Of these, Panamito and Canario are commercial varieties likely introduced from the coastal region. These are predominantly cultivated in higher-altitude zones by migrant farmers from the coast and highlands. The remaining varieties are traditional landraces distributed across both the lowland and upland Amazon. The most common variety collected was *Panamito*, representing 15.4% of all accessions. This was followed by local landraces such as *Waska* (12.4%), *Ashpa* (11.8%), and *Pinto* (6.5%).

**Table 2.** Nominal variability of common bean from Peruvian Amazonia.

| Common_name | Number | Frequency | Percentage (%) |
| --- | --- | --- | --- |
| Panamito | 143 | 0.15409483 | 15.41 |
| Waska | 115 | 0.12392241 | 12.39 |
| Ashpa | 109 | 0.1174569 | 11.75 |
| Pinto | 60 | 0.06465517 | 6.47 |
| Canario | 36 | 0.0387931 | 3.88 |
| Poroto | 36 | 0.0387931 | 3.88 |
| Ñuña | 34 | 0.03663793 | 3.66 |
| Shingo | 31 | 0.03340517 | 3.34 |
| Manteca | 27 | 0.02909483 | 2.91 |
| Awisho | 23 | 0.02478448 | 2.48 |
| Ojo de cabra | 22 | 0.0237069 | 2.37 |
| Cambio-90 | 20 | 0.02155172 | 2.16 |
| Chaucha | 20 | 0.02155172 | 2.16 |
| Norteño | 20 | 0.02155172 | 2.16 |
| Cápsula | 14 | 0.01508621 | 1.51 |
| Vaca paleta | 14 | 0.01508621 | 1.51 |
| Chiclayo | 11 | 0.01185345 | 1.19 |
| Pozuzo | 11 | 0.01185345 | 1.19 |
| Ashpilla | 10 | 0.01077586 | 1.08 |
| Garrapata | 10 | 0.01077586 | 1.08 |
| Mantequilla | 10 | 0.01077586 | 1.08 |
| Bayo | 9 | 0.00969828 | 0.97 |
| Caballero | 9 | 0.00969828 | 0.97 |
| Ninaporoto | 9 | 0.00969828 | 0.97 |
| San Pablino | 9 | 0.00969828 | 0.97 |
| Ucayalino | 9 | 0.00969828 | 0.97 |
| 60 días | 8 | 0.00862069 | 0.86 |
| Chuncho | 7 | 0.0075431 | 0.75 |
| Pintado | 6 | 0.00646552 | 0.65 |
| Vaquita | 6 | 0.00646552 | 0.65 |
| leche | 5 | 0.00538793 | 0.54 |
| frijol local | 5 | 0.00538793 | 0.54 |
| Cuarentón | 4 | 0.00431034 | 0.43 |
| Dark | 4 | 0.00431034 | 0.43 |
| Reclina | 4 | 0.00431034 | 0.43 |
| Blanco | 3 | 0.00323276 | 0.32 |
| Chaoca | 3 | 0.00323276 | 0.32 |
| Nube | 3 | 0.00323276 | 0.32 |
| Panamo | 3 | 0.00323276 | 0.32 |
| Burra | 2 | 0.00215517 | 0.22 |
| Caraota | 2 | 0.00215517 | 0.22 |
| Chinito | 2 | 0.00215517 | 0.22 |
| Cumbeño | 2 | 0.00215517 | 0.22 |
| Frejol de suelo | 2 | 0.00215517 | 0.22 |
| Pavita | 2 | 0.00215517 | 0.22 |
| Región | 2 | 0.00215517 | 0.22 |
| Sangre de toro | 2 | 0.00215517 | 0.22 |
| Amarillo | 1 | 0.00107759 | 0.11 |
| Arverja | 1 | 0.00107759 | 0.11 |
| Bayito | 1 | 0.00107759 | 0.11 |
| Bola | 1 | 0.00107759 | 0.11 |
| Camote | 1 | 0.00107759 | 0.11 |
| Canarillo | 1 | 0.00107759 | 0.11 |
| Comandante | 1 | 0.00107759 | 0.11 |
| Grande | 1 | 0.00107759 | 0.11 |
| Huallaguino | 1 | 0.00107759 | 0.11 |
| Huanuqueño | 1 | 0.00107759 | 0.11 |
| Huevo de pajarito | 1 | 0.00107759 | 0.11 |
| Lanche | 1 | 0.00107759 | 0.11 |
| Lentreja | 1 | 0.00107759 | 0.11 |
| Limón | 1 | 0.00107759 | 0.11 |
| Mani | 1 | 0.00107759 | 0.11 |
| Mocho | 1 | 0.00107759 | 0.11 |
| Nina panamito | 1 | 0.00107759 | 0.11 |
| Riñón | 1 | 0.00107759 | 0.11 |
| Rojo | 1 | 0.00107759 | 0.11 |
| Rundo | 1 | 0.00107759 | 0.11 |
| Seda | 1 | 0.00107759 | 0.11 |
| Tabaquero | 1 | 0.00107759 | 0.11 |
| Trujillano | 1 | 0.00107759 | 0.11 |
| Tumba-maíz | 1 | 0.00107759 | 0.11 |
| Vainita | 1 | 0.00107759 | 0.11 |
| Wayruro | 1 | 0.00107759 | 0.11 |
| Total | 928 | 1 | 100 |

### Morphological variability

We describe the characterization of 476 accessions (from 667 sowed), the rest of 191 accessions did not progress in the field trial. Based on qualitative and quantitative morphological descriptors, we estimated the variability of the collected accessions. Substantial phenotypic variability was observed among the accessions, particularly in traits such as seed color and growth habit, reflecting both natural selection and farmer-driven selection (Figure 2).

**Figure 2.**
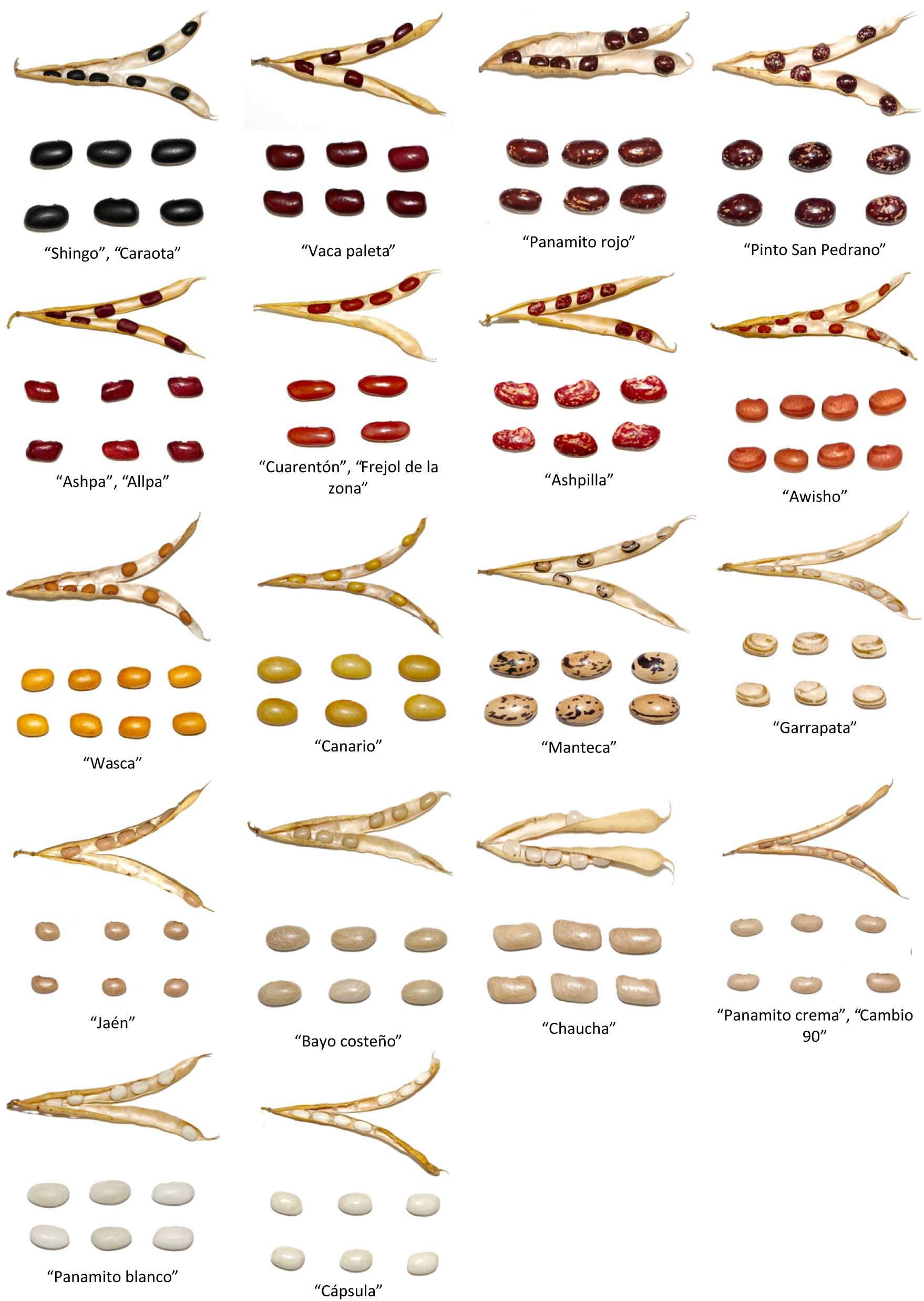
Morphological variability of common bean from Peruvian Amazonia, showing different form, size, color of seed and its ripe pod.

Boxplot for quantitative traits shows high variability, flowering time varying from 32 to 75 days, in the case of pod length the range was from 6.5 to 17.3 cm and locules per pod ranged from 2 to 11; which are very important source for common bean plant breeders (Figure 3). Also, the histogram of qualitative traits shows variability in the color of flower, pod and seed, also different size and shapes of the pods and seeds (Figure 4), which is a great variability for different use of the germplasm.

**Figure 3.**
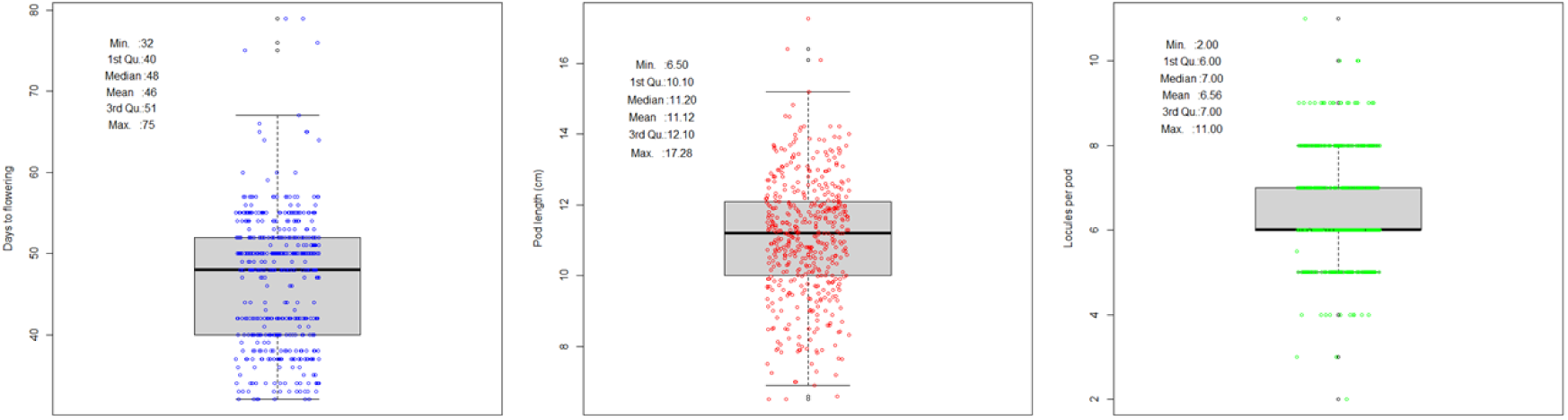
Boxplot of quantitative descriptors on 496 common bean accessions.

**Figure 4.**
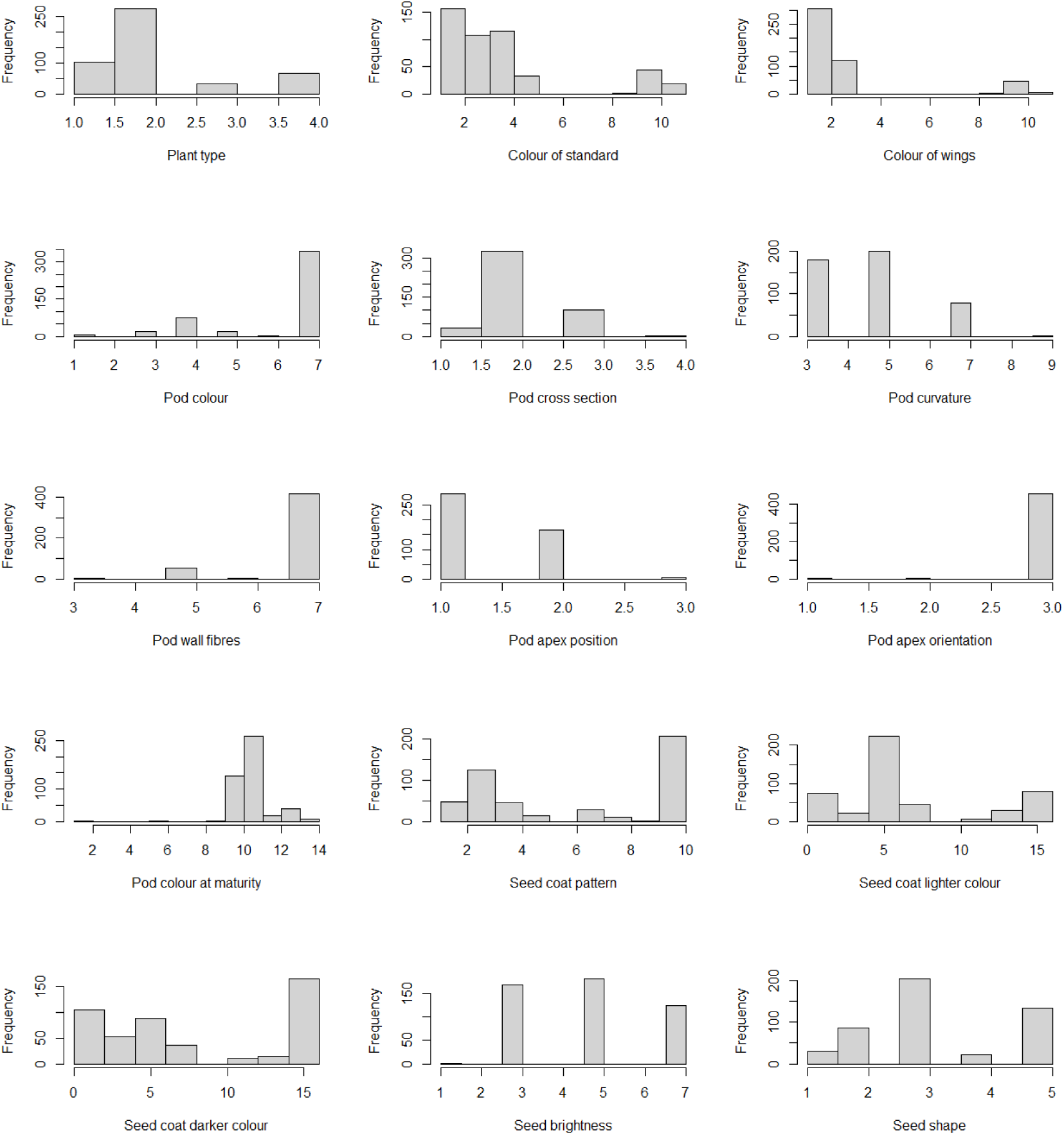
Histogram of qualitative descriptors on 496 common bean accessions.

**Figure 5.**
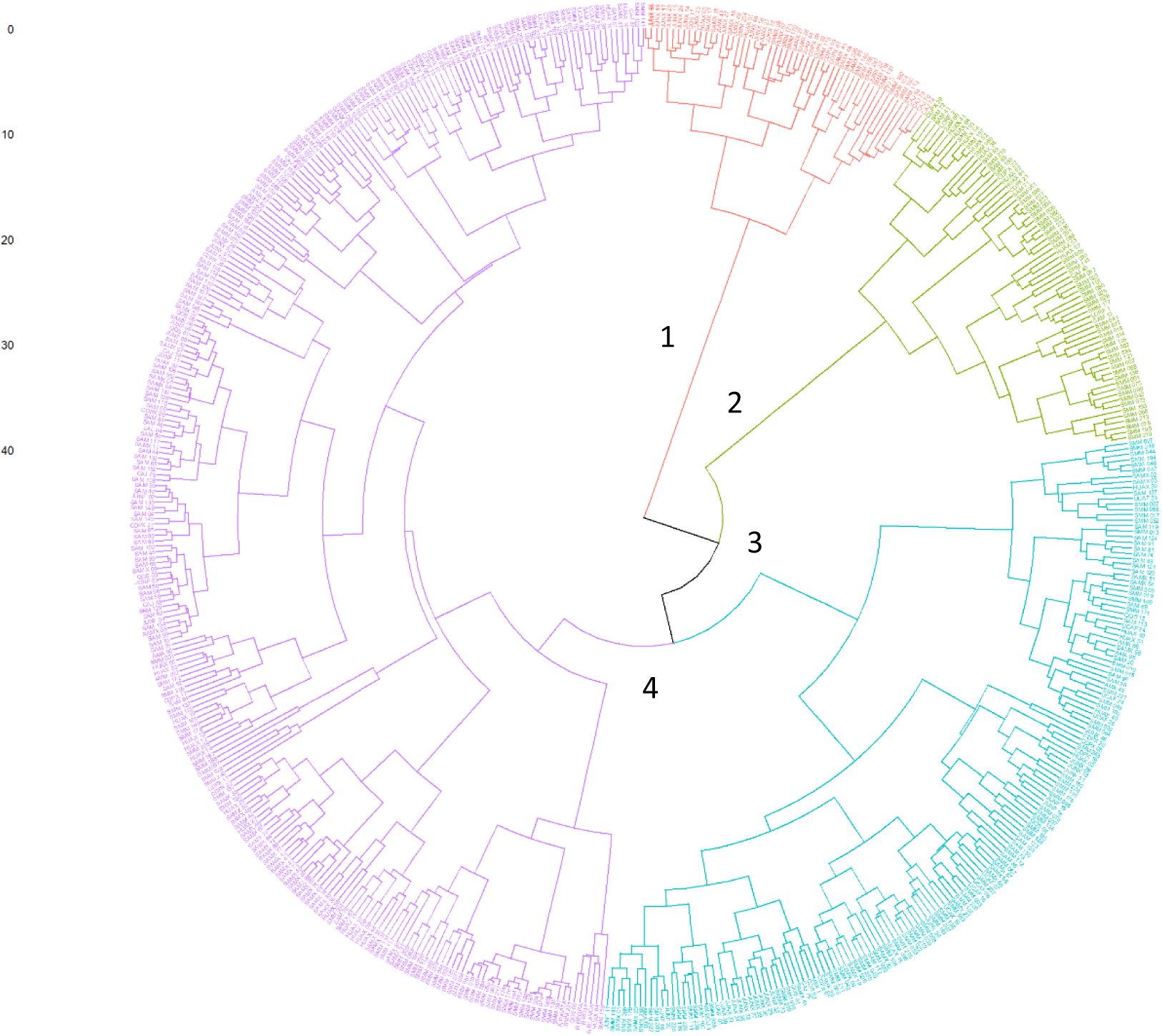
Tree of 476 common bean accessions according to 19 morphological descriptors (group 1 = red, group 2 = green, group 3 = Blue, and group 4 = pink).

### Relationships and variability

Cluster analysis based on morphological traits grouped the accessions into four major clusters (Figure 3). These clusters were not strictly associated with geographic origin; however, some accessions—such as *Awisho* and *Ashpa*,—tended to group together and were commonly cultivated in lowland areas. This suggests possible morphological differentiation based on ecological zones, particularly altitude-related environmental conditions.

Cluster 1 are grouping accessions from San Martin, Huánuco and Cerro de Pasco regions, the type of plants are mostly indeterminate climbing or indeterminate semi-vine/prostrate, white standard color, white wings, green pods, the pod cross-section round elliptic, pod Slightly curved, in average present 8 locules per pod, excessive shattering of pod wall fibers, marginal pod apex position, downwards pod apex orientation, the pods color at maturity are golden yellow, the main seed coat color is yellow, without secondary coat pattern, and the seed shape is varying through to the all states of this descriptor, from round to truncate fastigiate.

Cluster 2 are grouping accessions from San Martin, Huánuco, Cerro de Pasco and Junin regions, the type of plants are indeterminate bushy, the standard color is variable from white to purple, white wings, normal green pods, Pear shaped pod cross-section, pod curvature variable from Straight to Recurving, in average present 5.9 locules per pod, excessive shattering of pod wall fibers, this group present marginal or non-marginal pod apex position, in the pod apex orientation present two estates straight and downwards, the pod color at maturity varying in different degree of yellow, the Seed coat pattern is varying through the different states (constant mottled, striped, rhomboid spotted, speckled, broad striped; 9 Spotted bicolor), the seed coat color in mainly pale-cream and pure white, with variable secondary coat pattern as maroon, red and pink, the seed brightness are shiny and medium and the seed shape is varying through to the states as oval, cuboid, kidney shaped and truncate fastigiate.

Cluster 3 are grouping accessions from San Martin, Huanuco, Cerro de Pasco and Junin regions, the state differentiation for this group it was found in the following descriptors: Color of wings whit white and lilac states, pod wall fibers the state mostly was excessive shattering and followed by leathery podded and on pod color at maturity varying in different degree of yellow. However, for thew rest of descriptors there is no clear of the state differentiations to associate exclusive to this group.

Cluster 4 are grouping accessions from San Martin, Huanuco, Cerro de Pasco and Junin regions, the type of plants were indeterminate bushy and determinate bushy, the standard color was different deep of lilac and purple, the wings were purple, normal green pods, mostly pear shapped pod cross-section, pod curvature Straight, pod wall fibers excessive shattering, non-marginal pod apex position, the pod color at maturity was clear yellow. For the rest of descriptors there is no a clear differentiation for this group, it seems a mixed germplasm.

The first two Principal Component (PC) explain 16.9% and 11.2% respectively, and the cumulative explain 28.1 % (Figure 6), this result means there is a high variability, because to explain others 71.9% of the variability it is needed more PC. The PCA-biplot shows four groups, similar than cluster analysis, only group one and group four are clearly differentiated according to the morphological traits. The group one shows late flowering time (around 50 days), Excessive shattering of pod wall fibers, kidney shaped and truncate fastigiate of seed shape, high number of locules per pod (around 8); contrary the group four early plants (around 38 days), strongly contracting of pod wall fibers, round and Oval of seed shape, low number of locules per pod (around 5.6). For the groups two and three there is no clearly separated, it seems admixture between them.

**Figure 6.**
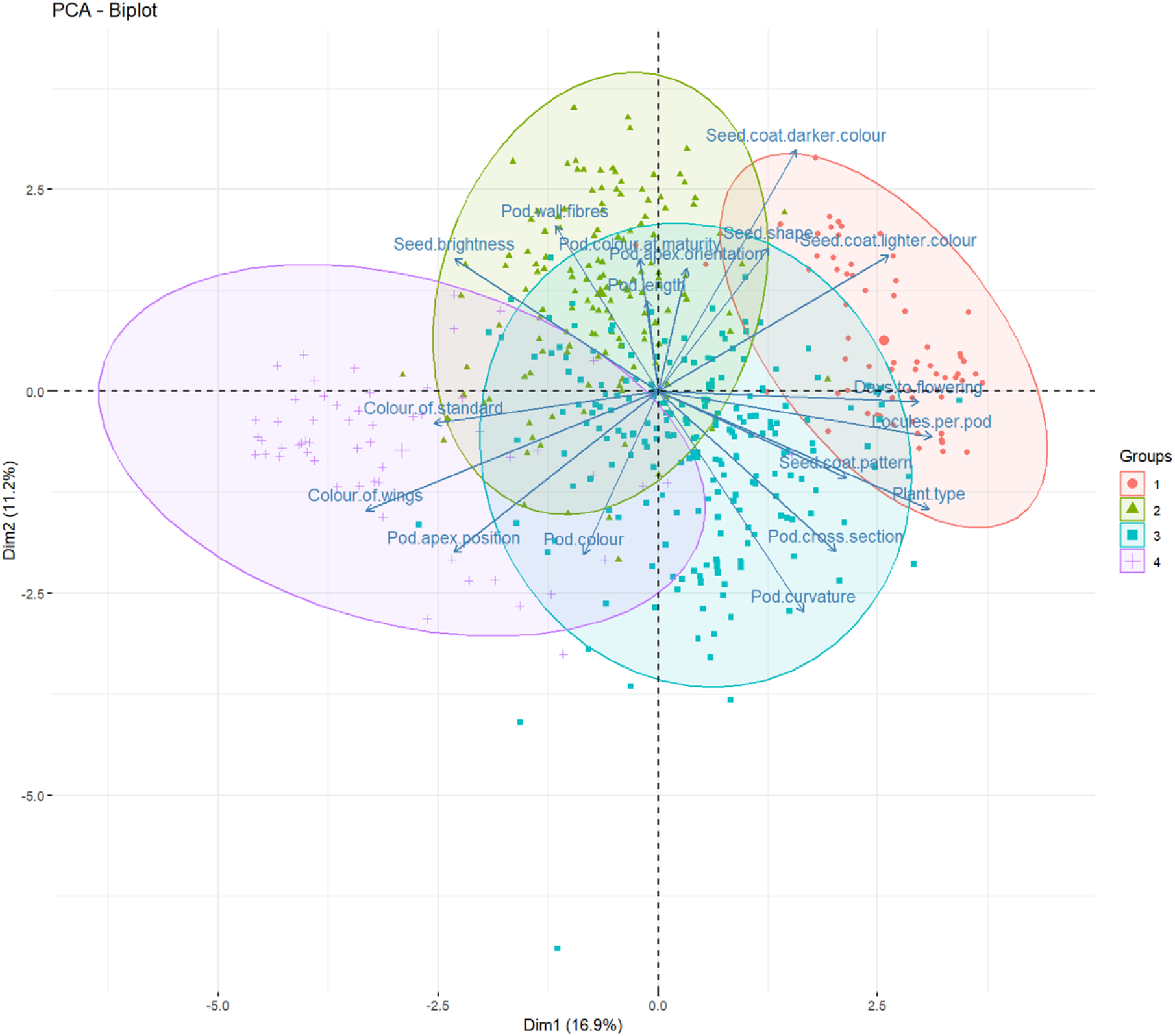
The two first PC biplot of 476 common bean accessions according to 18 morphological descriptors.

Multiple Correspondence Analysis (MCA) taking into consideration the subgroups found according to SNP data, in overall shows that the subgroups do not have exclusive discriminating qualitative descriptors which differentiate them from each other (Figure 7). There is only an exception with subgroup 1.2, which has a lilac flower color (standard and wings) and a light red pod color at physiological maturity and a carmine seed coat color. This subgroup belonging to the Andean gene pool.

**Figure 7.**
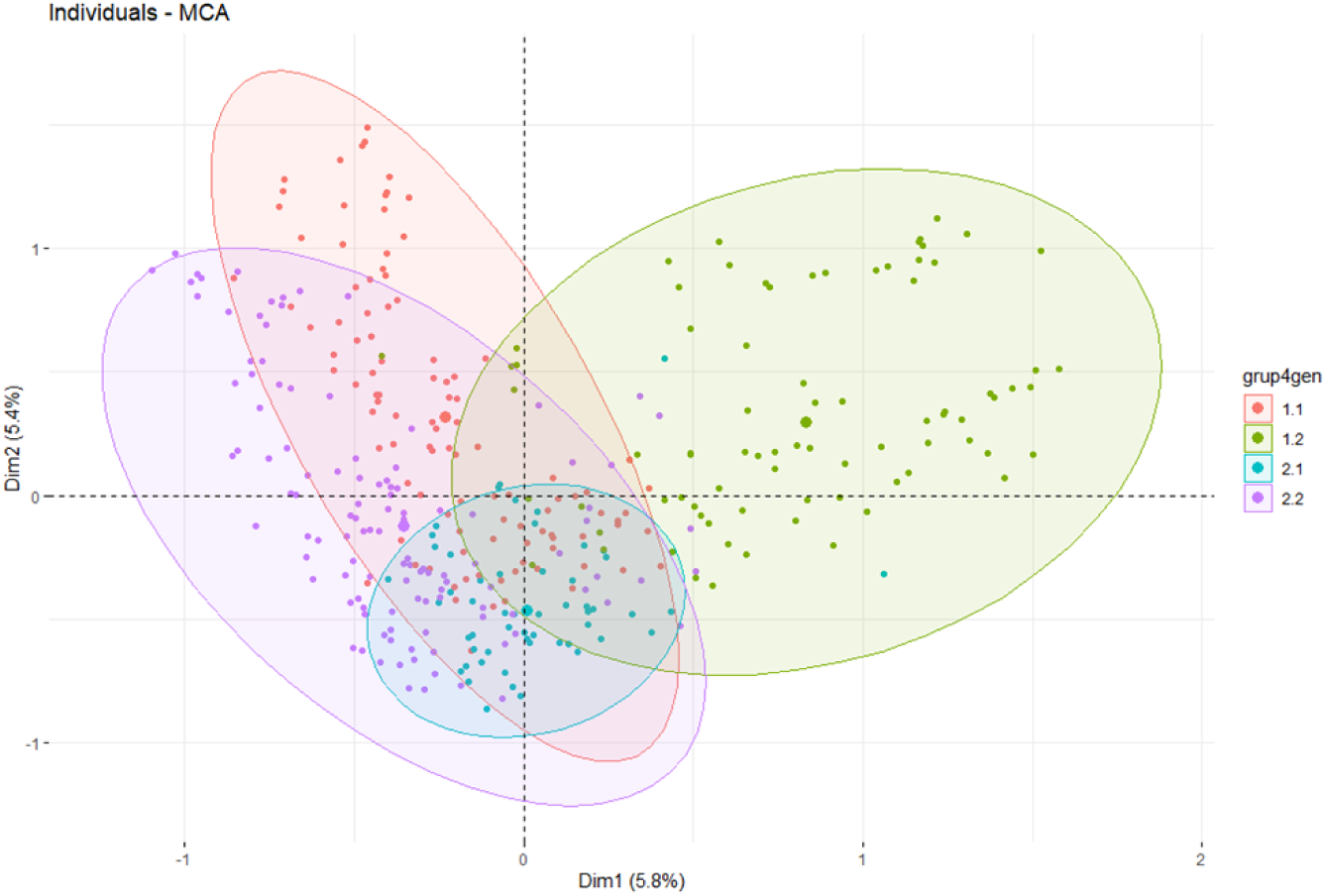
Multiple Correspondence Analysis (MCA) of 476 accessions with 15 qualitative descriptors.

### SNP marker variation and distribution

From the initial set of 665 common bean accessions, 647 presented less than 30% missing data and were retained for downstream analyses. SNP markers were filtered to retain only those with a call rate of at least 50% and a minor allele frequency (MAF) of 1% or higher, yielding a filtered dataset of 24,283 high-quality SNPs (Figure 8, Table 3). Alignment to the common bean YP4 reference genome revealed that 23,050 SNPs (94.9%) were successfully mapped to chromosome positions.

**Figure 8.**
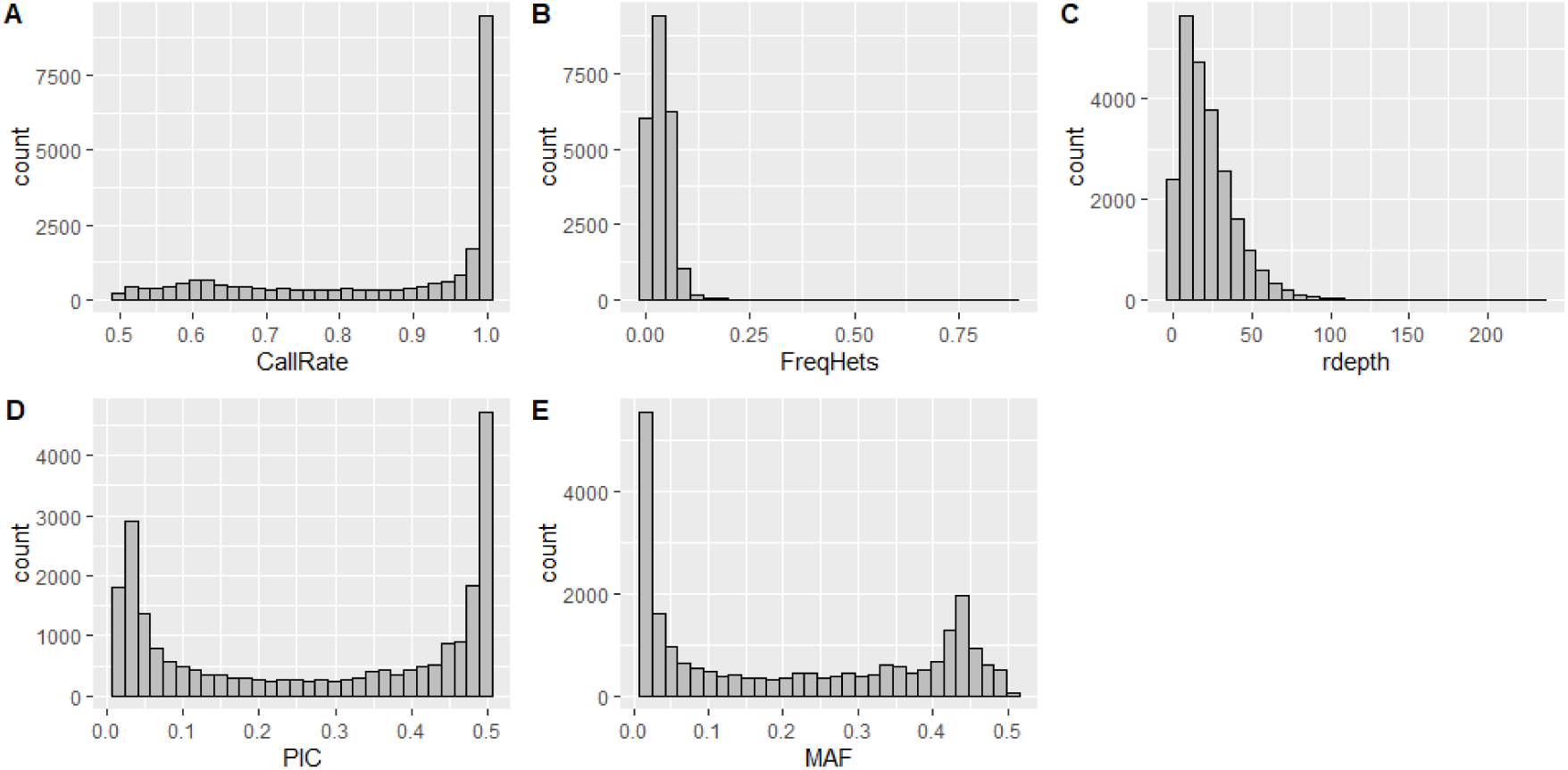
Histogram distribution of SNP marker parameters A) Callrate, B) Frequency of heterozygous genotypes, C) read depth, D) Polymorphic Index Content (PIC), and E) minimum allele frequency (MAF) on the filtered set of 23050 SNP markers.

**Table 3.** Single nucleotide polymorphism (SNP) call rate report pre and post filtering.

|  | Reporting Call Rate by |  | Call Rate post filtering |  |
| --- | --- | --- | --- | --- |
|  | Individual | Locus | Individual | Locus |
| No. of loci | 80079 | 80079 | 23050 | 23050 |
| No. of individuals | 665 | 665 | 647 | 647 |
| Minimum | 0.3658887 | 0.344361 | 0.500773 | 0.7078525 |
| 1st quartile | 0.8411069 | 0.701504 | 0.714065 | 0.8309761 |
| Median | 0.8577155 | 0.966917 | 0.967543 | 0.8640781 |
| Mean | 0.8507188 | 0.8507188 | 0.8623075 | 0.8623075 |
| 3r quartile | 0.8785449 | 0.989474 | 0.998454 | 0.8944252 |
| Maximum | 0.9769603 | 1 | 1 | 0.9823861 |
| Missing Rate Overall | 0.1493 | 0.1493 | 0.1377 | 0.1377 |

For SilicoDArT markers, the applied filter of Call rate ≥ 90% and MAF of ≥ 1% produced a set of 29,865 markers. Additionally, alignment to YP4 genome reference revealed that 22,594 SilicoDArT markers (75.7%) were successfully mapped to chromosome positions. Results of statistical analysis were very similar than obtained with SNP marker, which is shown as supplementary material (Figures 13, 14, 14 and 15).

### Genetic relationships and Population Structure

Clustering analysis using the neighbor-joining algorithm grouped the accessions into two main groups, reflecting underlying genetic relationships (Figure 9). The first group consists primarily of accessions from the Peruvian Amazon, plus samples added as an outgroup from Andean origin (Peruvian highlands and coast, and accessions from CIAT). The second group consists of accessions from the Peruvian Amazon plus samples added as an outgroup from Mesoamerica origin (material from CIAT, and a sample from Mexico). This indicates that the first group belong to the Andean gene pool, and the second one to the Mesoamerican gene pool.

**Figure 9.**
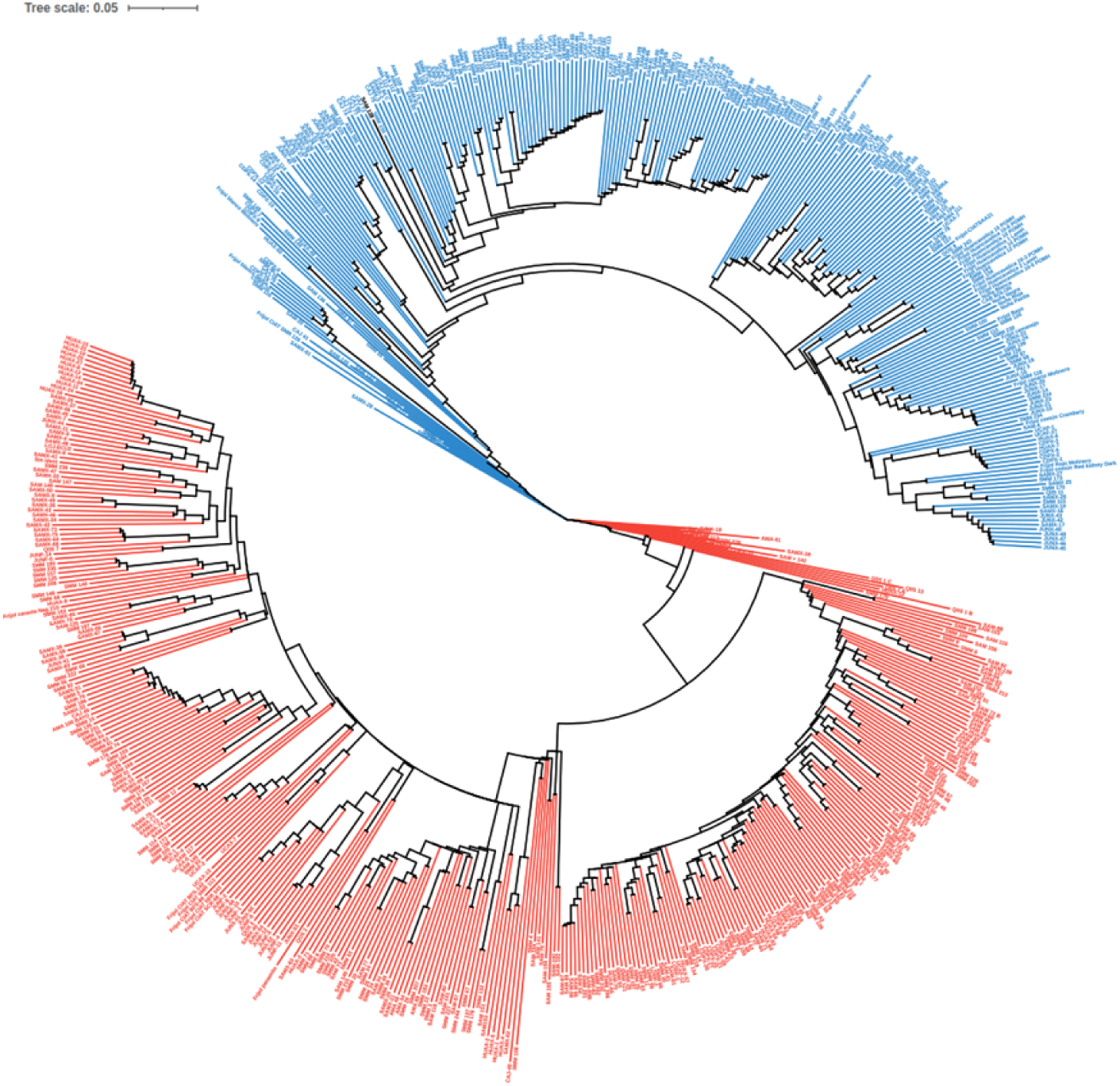
Dendrogram of 647 accessions of common bean, according to the 23,050 DarTseq single nucleotide polymorphism (SNP) markers.

Principal Coordinate Analysis (PCoA) further supported these findings, with the first three axes explaining 62.62%, 8.97% and 2.77% of the total variance and clearly separating the two main groups within the collection (Figure 10). Evanno plot showing the strongest signal of population structure at K=2, which is supported by cluster analysis identified two major genetic groups and PcoA, correlating with Andean and Mesoamerican gene pool origin and likely according to its domestication and adaptation history (Figure 11). Within these groups, two subgroups are also observed, as shown in Figure 10, with k = 4. These subgroups are clustering accordingly its common name, which is a clue to build adaptation process and cropping history of common bean in the Amazonia (Figure 12).

**Figure 10.**
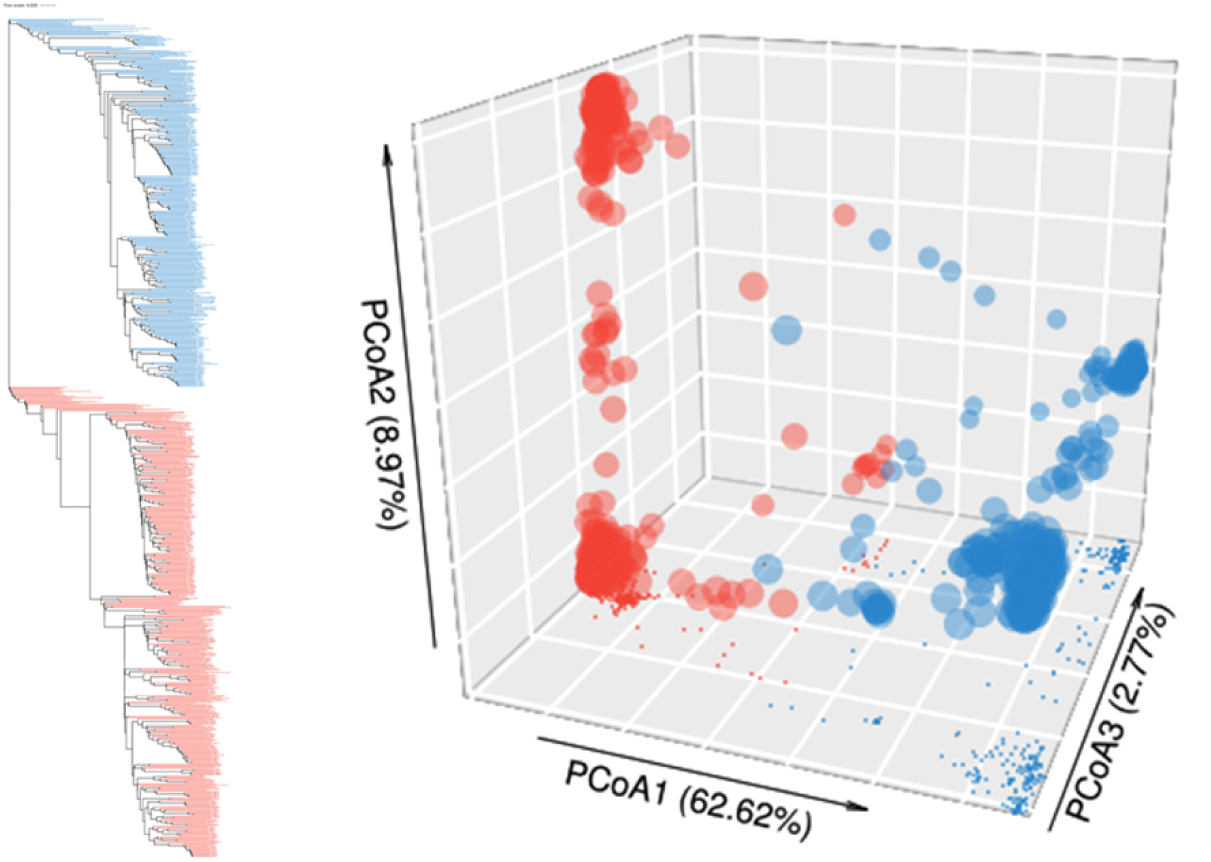
PcoA of 647 accessions of common bean according to the DarTseq single nucleotide polymorphism (SNP) marker data.

**Figure 11.**
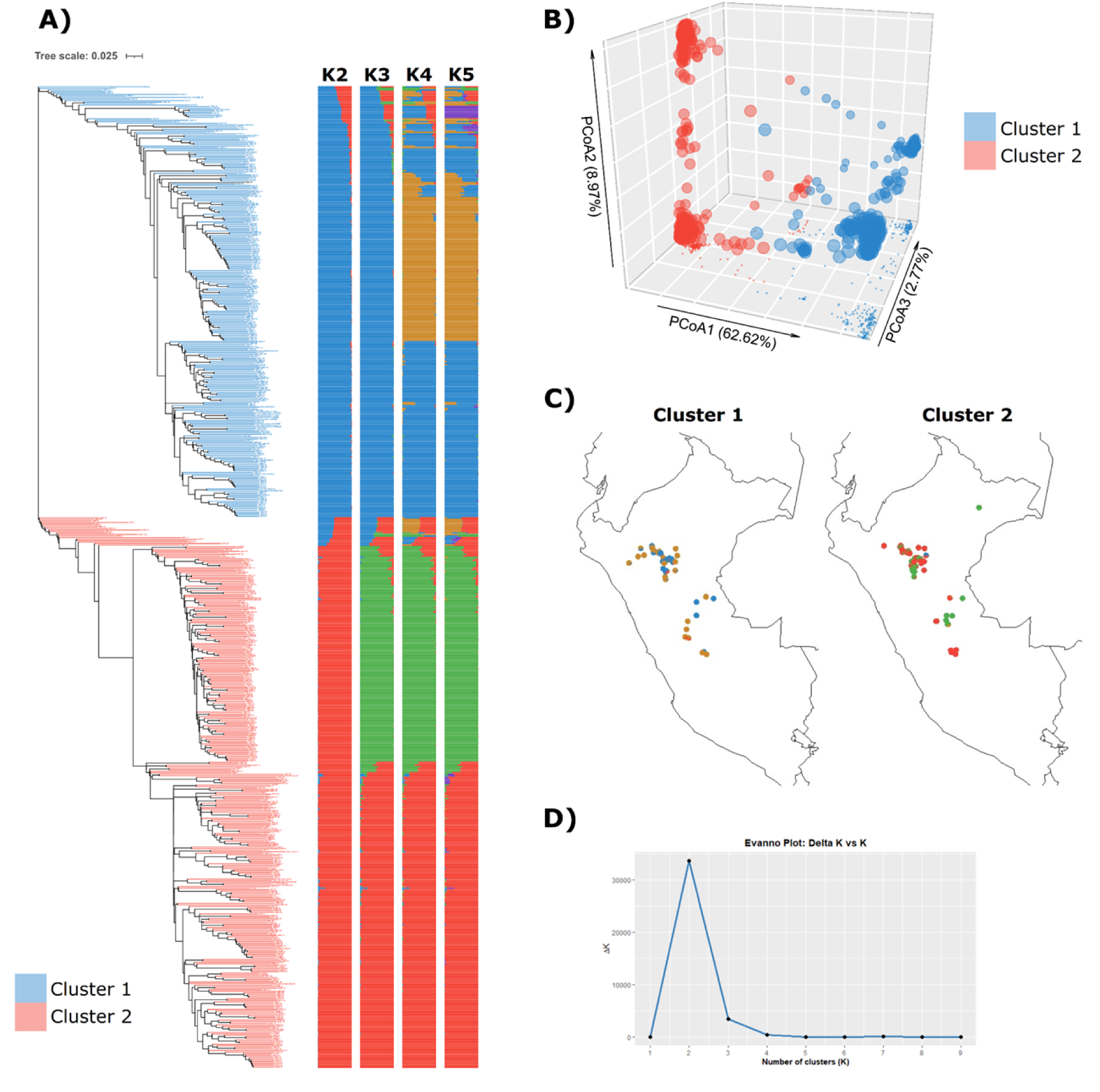
Genetic structure of 647 common bean accessions with SNPs 23,050 markers. A) Neighbor joining tree showing two mainclusters, ancestry proportions from STRUCTURE analysis from K2 to K5 (left to right). B) Principal Coordinate Analysis colored according to the two main clusters of the NJ tree. C) Pie chart of the ancestry proportions of K=4 for each accession with geographic coordinates. D) Evanno plot showing the strongest signal of population structure at K=2.

**Figure 12.**
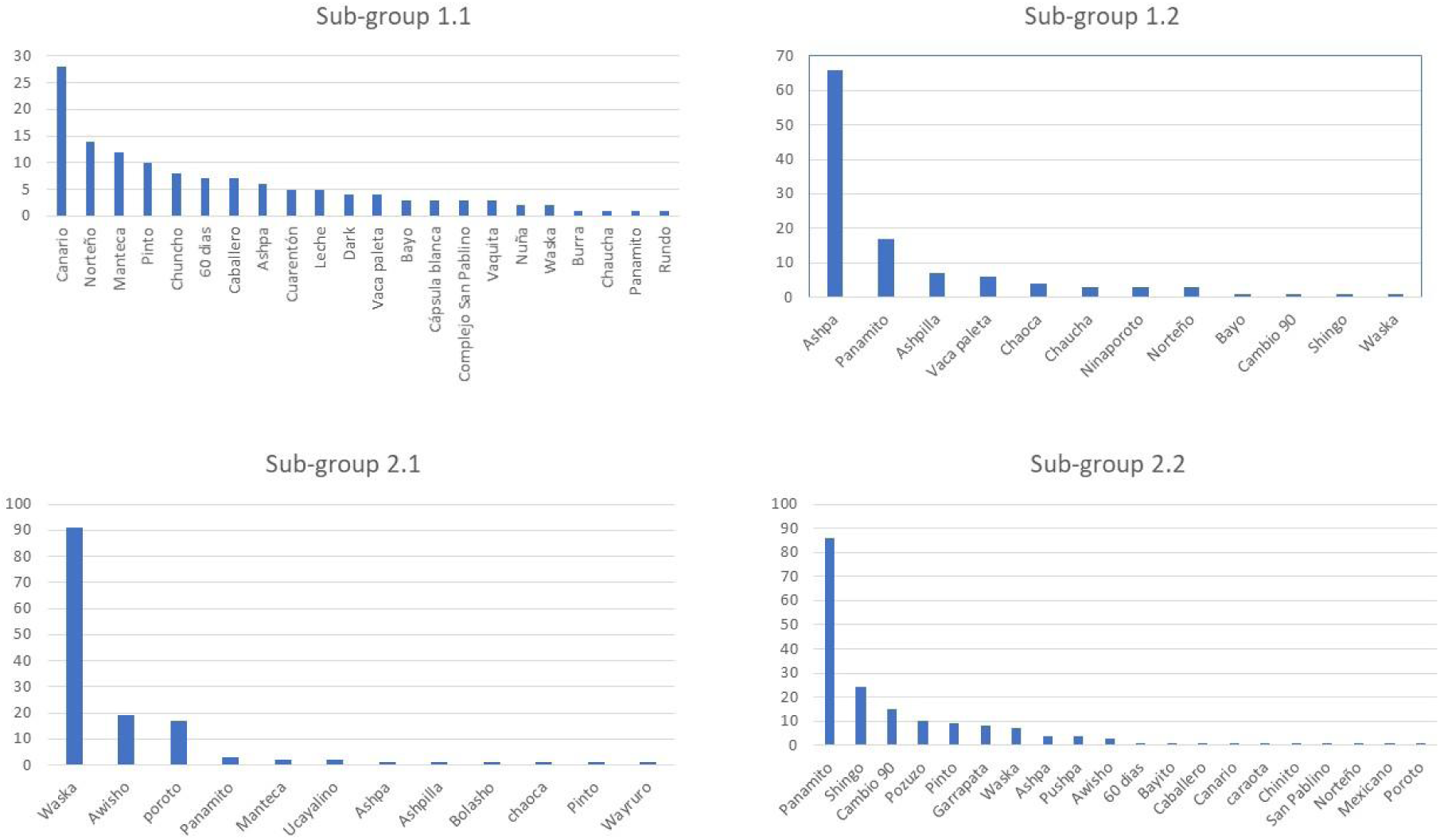
Number of common names distribution in each subgroup of common bean from Amazonia, Peru.

The subgroups corresponding to the first group have been designated as subgroup 1.1 and subgroup 1.2. Subgroup 1.1 consists of recently collected samples and samples from the mountains (popping beans-nuñas, beans) and the coast, which were added as separate groups to observe the genetic relationships among Peruvian beans. This subgroup predominantly contains beans with the following common names: Canario, Norteño, Manteca, Pinto, Chuncho, 60 días, and Caballero. While, subgroup 1.2 consists of all recently collected samples from the Peruvian Amazon; this subgroup predominantly contains beans with the following common names: Ashpa, Panamito, Ashpilla, and Vaca paleta (Figure 12).

The subgroups corresponding to the second group have been named as subgroup 2.1 and subgroup 2.2. Subgroup 2.1 consists of all recently collected samples from the Peruvian Amazon; this subgroup predominantly contains common beans cultivated with the following common names: Waska, Awisho, and poroto. In contrast, subgroup 2.2 consists of recently collected samples from Peruvian amazon and samples added as an outgroup from Mesoamerica origin; this subgroup predominantly contains beans with the following common names: Panamito, Shingo, Cambio 90, Pozuzo, Pinto, and Garrapata (Figure 12).

### Genetic Diversity (α and β) using “q” profiles

Using the two main groups identified in the Peruvian Amazon common bean panel, we quantified within-group (alpha, α) and between-group (beta, β) diversity across Hill numbers for q = 0, 1, and 2 based on the 23,050 SNP markers. The results are shown in the table 4.

**Table 4.** Hill number of α (intrapopulation) and β (interpopulation) diversity indices according to the 23,050 loci SNP and two groups of common beans from Amazonia, Peru.

| Hill numbers (D) | Measures | Alpha diversity ($\alpha$) | | Beta diversity ($\beta$) |
| --- | --- | --- | --- | --- |
|  |  | Group1 | Group2 | Differentiation of group 1-group 2 |
| q=0 (allele richness) | Potential richness | 1.97 | 1.95 | ~1.04 |
| q=1 (Shannon index) | Frequency balance | 1.31 | 1.31 | ~1.2 |
| q=2 (expected heterozygosity) | Effective heterozygosity (weight on common alleles) | 1.22 | 1.23 | ~1.4-1.5 |

For q = 0 (allelic richness), α was 1.97 in Group1 and 1.95 in Group2, which is expected in biallelic SNPs, while β was 1.04, indicating that both groups harbor essentially the same set of allelic states with virtually no private alleles unique to either group. At q = 1 (Shannon diversity), α was identical in both groups (1.31) and β increased to 1.20, which indicates that although within-group allele-frequency balance is similar, the groups begin to differ in how common those alleles are between populations. At q = 2 (Gini–Simpson/expected heterozygosity), α was 1.22 in Group1 and 1.23 in Group2, whereas β rose to 1.4–1.5, indicating a clearer differentiation driven primarily by shifts in the frequencies of common alleles.

Taken together, these q-profiles show minimal presence/absence differentiation but progressively stronger frequency-based divergence from q = 0 to q = 2, indicating that the genetic structuring between groups is explained more by differences in the prevalence of shared alleles than by unique allelic repertoires.

Using the same 23,050 SNPs, we extended the q-profile analysis to four populations defined by previous cluster and STRUCTURE analysis. The results of the analysis in the four subgroups are shown in Table 5.

**Table 5.** Hill number of α (intrapopulation) and β (interpopulation) diversity indices according to the 23,050 loci SNP and four subgroups of common beans from Amazonia, Peru.

| Subgroups | q=0 | q=1 | q=2 | Diversity |
| --- | --- | --- | --- | --- |
| Subgroup1_1 | 1.93 | 1.31 | 1.22 | $\alpha$ |
| Subgroup1_2 | 1.89 | 1.25 | 1.16 |  |
| Subgroup2_1 | 1.82 | 1.16 | 1.1 |  |
| Subgroup2_2 | 1.94 | 1.3 | 1.21 |  |
| Comparison subgroup1_1<br>vs subgroup1_2 | 1.06 | 1.24 | ~1.29 | $\beta$ |
| Comparison subgroup2_1<br>vs subgroup2_2 | 1.07 | 1.29 | ~1.32 |  |

For q = 0, α ranged narrowly from 1.82 to 1.94 (see Table 5), indicating that all populations harbor nearly the full set of biallelic states with only minor differences in allelic richness. At q = 1, α values differentiated the populations more clearly (Subgroup1_1 = 1.31; Subgroup1_2 = 1.25; Subgroup2_1 = 1.16; Subgroup2_2 = 1.30), which indicates varying degrees of allele frequency balance: Subgroup1_1 and Subgroup2_2 exhibit more even allele frequencies, whereas Subgroup2_1 shows greater skew toward common alleles. At q = 2, the ranking persisted (Subgroup2_1 = 1.10; Subgroup1_2 = 1.16; Subgroup2_2 = 1.21; Subgroup1_1 = 1.22), indicating that differences among populations are driven primarily by the frequencies of common alleles.

Taken together, the four-way comparison shows minimal differences in allelic repertoires (q = 0) but increasing divergence at q = 1 and q = 2, which indicates that genetic differentiation among these subpopulations is explained less by presence/absence of allelic states and more by shifts in the prevalence and balance of shared alleles.

Comparing the two subgroups within each main group we observe little presence/absence differentiation but a clear frequency-based divergence. For subgroup1_1 versus subgroup1_2, β increased from 1.06 at q = 0 to 1.24 at q = 1 and ∼1.29 at q = 2, which indicates that while both subgroups share essentially the same allelic states, they differ modestly to moderately in allele-frequency balance, with the strongest contrast driven by common alleles. A similar behavior is obseved for subgroup2_1 versus subgroup2_2 (Table 5).

### Population Heterozygosity

The parameters of diversity using heterozygosity, according to the 23,050 loci SNP, two-groups and four subgroups of common beans are shown in the table 6.

**Table 6.** Genetic diversity parameters within groups and subgroups with the 23,050 loci SNP of common beans from Amazonia, Peru.

| Groups | n.Ind (ef.) | n.Loc | polyLoc | monoLoc | Ho (±SD) | He (±SD) | uHe (±SD) | FIS (±SD) |
| --- | --- | --- | --- | --- | --- | --- | --- | --- |
| Group_1 | ~240 | 23,050 | 22,281 | 769 | 0.041 (0.056) | 0.148 (0.145) | 0.148 (0.145) | 0.523 (0.364) |
| Group_2 | ~318 | 23,050 | 21,846 | 1,204 | 0.035 (0.051) | 0.147 (0.165) | 0.147 (0.165) | 0.540 (0.365) |
| Subgroup1_1 | ~136 | 23,050 | 21,511 | 1,539 | 0.032 (0.054) | 0.145 (0.148) | 0.146 (0.149) | 0.576 (0.384) |
| Subgroup1_2 | ~104 | 23,050 | 20,444 | 2,606 | 0.053 (0.068) | 0.115 (0.124) | 0.116 (0.125) | 0.348 (0.380) |
| Subgroup2_1 | ~131 | 23,050 | 18,877 | 4,173 | 0.029 (0.058) | 0.071 (0.114) | 0.071 (0.114) | 0.302 (0.380) |
| Subgroup2_2 | ~187 | 23,050 | 21,563 | 1,487 | 0.040 (0.055) | 0.139 (0.149) | 0.139 (0.149) | 0.525 (0.355) |

n.Ind (ef.) = effective sample size; n.Loc = number of loci; polyLoc = number of polymorphic loci; monoLoc = monomorphic loci; Ho (Observed heterozygosity; proportion of heterozygous loci per individual, average across the population); He (Expected heterozygosity, Nei 1978; expected heterozygosity under Hardy–Weinberg equilibrium); uHe (unbiased He, corrected for sample size); FIS (Inbreeding coefficient = $1 - Ho / uHe$ , positive values indicate a deficit of heterozygotes (possible inbreeding or structure) and negative values indicate an excess).

### Two-groups analysis

Group 1: where the effective sample size was ∼240 individuals, showing 22,281 polymorphic loci (almost all, 97% of the total). The Ho = 0.041, meaning that on average, ∼4.1% of the loci per individual are heterozygous. Then, the He/uHe = ∼0.148, under Hardy-Weinberg Equilibrium (HWE) shows around ∼14.8% of heterozygous loci would be expected. The FIS = 0.52 (inbreeding coefficient), indicated a severe heterozygote deficit. The population is less heterozygous than expected, which reflect inbreeding, that is a expected result in a self-pollinating populations as a common bean.

Group 2: The effective sample size was ∼318 individuals, showing 21,846 polymorphic loci (94.8% of the total). The Ho = 0.035, around ∼3.5% of loci are heterozygous on average, even lower than group 1. The proportion He/uHe= ∼0.147 is similar to group 1 in expected heterozygosity; and FIS= 0.54, also shows strong heterozygote deficit, very comparable to group 1. Comparing both Group 1 vs. Group 2, they show very similar He (∼0.15), indicating that the allele pool and frequencies are similar. Ho is lower in group 2 than group 1(0.035 vs. 0.041), suggesting that individuals in group 2 express less heterozygosity. The FIS is elevated in both populations (>0.5), reinforcing the idea of a generalized heterozygote deficit, which is expected for common bean populations. Finally, group2 has more monomorphic loci than group 1 (1,204 vs 769), which could reflect loss of variation or fixation at certain sites.

### Four subgroups analysis

The subgroup1_1 has effective size ∼136 individuals and 21,511 polymorphic loci (93% of the total). The Ho = 0.032, quite low compared to He = 0.145; and the FIS = 0.58, reflecting a strong heterozygote deficit.

The subgroup 1_2 has effective size ∼104 individuals and 20,444 polymorphic loci (88.7%), more monomorphic loci than subgroup1_1. The Ho = 0.053, higher than subgroup1_1, individuals show slightly more heterozygosity. However, the He = 0.115 was lower than subgroup1_1; and the FIS = 0.35, still positive but lower heterozygote deficit than in subgroup1_1.

Subgroup2_1 has effective size ∼131 individuals and 18,877 polymorphic loci (82%), the lowest of the four subgroups. Also, the Ho = 0.029 and He = 0.071 were the lowest of all subgroups, indicating a loss of diversity; and the FIS = 0.30 showed a moderate heterozygote deficiency.

Subgroup2_2 has effective size ∼187 individuals and 21,563 polymorphic loci (93.5%), comparable to subgroup1_1. The Ho = 0.040 showed an intermediate value between subgroup1_1 and subgroup1_2; and the He = 0.139 was intermediate too; and FIS = 0.53, indicating a severe heterozygote deficiency.

### Comparisons between subgroups within the group

Group_1: the subgroup1_1 has a higher He (0.145) but a lower Ho, resulting in a very high FIS (0.58). While, the subgroup1_2 has a lower He (0.115) but a relatively higher Ho (0.053), which reduce the deficit of heterozygosity (FIS = 0.35). The subgroup1_1 appears more structured as an self-pollinating population, while subgroup1_2 maintains more heterozygotes despite lower overall diversity.

Group 2: the subgroup2_1 has the lowest diversity among the four subgroups (Ho = 0.029; He = 0.071). However, the subgroup2_2 recovers diversity (He = 0.139), but its FIS is still high (0.53). Within Group_2, there is a clear inequality; subgroup2_1 present loss of diversity; and subgroup2_2 is more diverse but lacks heterozygotes.

In all four subgroups, there is a clear disconnection between Ho and He, all subgroups have fewer heterozygotes than expected under Hardy–Weinberg equilibrium. Also, the positive and high FIS (0.30–0.58) in all subgroups suggest inbreeding, internal structure favoring homozygotes, which is expected in common bean.

## Discussion

### Morphologic variability

The morphologic diversity observed in term of type of plant, flowering time, color, size, form of pods and seeds confirms that Amazonian *P. vulgaris* represents an important genetic reservoir. The presence of distinct genetic clusters suggests that both environmental heterogeneity and farmer practices shape phenotypic variability. Similar levels of morphological diversity have been reported in Brazil [42], Mexico [43], Spain [44], Nigeria [45], Pakistan [46], Ethiopia [47]. In Peru, however, most previous studies have focused on highland germplasm, using small sample size [23] [22] [21] [48] and lately sample from Amazonia [25]; which are not representative to allow a comprehensive estimation of common bean variability. Our results, show that much of the morphological diversity is conserved on farms, where farmers save, select, and replant seeds year after year, ensuring both autonomy and household income [49] [50]. In addition, frequent seed acquisition by farmers from traditional markets further promotes gene flow and admixture, particularly relevant under tropical conditions where high temperature and humidity shorten seed longevity, and its storage challenges; causes rapid loss of viability. Together, these practices contribute to the maintenance and renewal of agrobiodiversity in the Peruvian Amazon. It is relevant to highlight the farmer behaviors, who collet new material and sow all together with own local landraces. Then, the agroecological mosaic, coupled with traditional knowledge, has led to the maintenance of a broad range of landraces with varying morphological traits. A large portion of this agrobiodiversity is still conserved *in situ*, within Indigenous and local farming systems. Farmers contribute to the conservation and evolution of these varieties by using and exchanging seeds year by year [51] [52]. Additionally, field observations and interviews revealed that many farmers regularly acquire seeds from traditional markets to replace planting stock each season. This behavior is likely influenced by tropical environmental constraints—such as high temperatures and humidity—which accelerate seed aging and reduce viability during storage. Thus, constant renewal of planting material becomes a necessary strategy. The morphological diversity recorded—particularly in traits like growth habit, seed color, and pod characteristics— supports the idea of local adaptation and cultural preferences influencing varietal selection. Morphological characterization, though often considered basic, remains a valuable and accessible tool for identifying distinct genotypes and understanding their agronomic potential and farmer preferences.

### Molecular diversity

SNP-based analyses, cluster and Principal Coordinate analysis (PcoA) revealed that the accessions were assigned into two distinct groups, corresponding to the Andean or Mesoamerican gene pools, 284 and 363 accessions, respectively (Figure 11). Of the Amazonia accessions, 56.1 % clustered with Mesoamerican pool and 43.9 % with the Andean gene pools. However, this tendency was largest in others common bean germplasm genetic diversity studies, e.g. in Brazil was reported, more than 70 % of Mesoamerican gene pool, [4]; [53]; and in Ethiopian germplasm, the 284 common bean accessions were roughly classified into Andean and Mesoamerican gene pools, assigning 188 (65.05%) accessions in the Mesoamerican group, while the remaining 96 (33.22%) accessions were classified in the Andean gene pool [28]. The population structure analysis (based on K = 2 groups) showed that the common bean accessions from Peruvian amazon were distributed between two gene pools, Andean and Mesoamerican with a low degree of admixture. By comparison, when considering K = 4, both the Andean and Mesoamerican gene pool was clustered in two subgroups, with low admixture.

This structure is consistent with findings from USDA core collections [54], where PCoA and NJ trees confirmed the Andean–Mesoamerican split and further subdivisions into ecogeographic races. These results indicate that Amazonian germplasm is integrated into the global diversity framework of *P. vulgaris* while retaining unique local substructure. However, in Brazilian germplasm genetic diversity studies in common bean were obtained more diversity in Mesoamerican gene pool using GBS analysis [53]. In overall, the genetic variation detected through DarTseq SNP analysis showed that, the clustering agrees with that obtained by the principal coordinate analysis and population structure analyses.

The high genetic variation detected through DArTseq SNP analysis complements the phenotypic findings and highlights the importance of both in situ and ex situ conservation. Many of these accessions, particularly those maintained only on-farm, are at risk of being lost due to changes in land use, market pressures, climate change, or generational shifts in farming practices. Comprehensive genetic characterization enables the identification of unique and underrepresented alleles, which can be prioritized for conservation and potentially incorporated into breeding programs aimed at improving resilience to biotic and abiotic stresses.

### Alpha (α) and Beta (β) Genetic Diversity

The Peruvian Amazonia common beans, according to the alpha-diversity estimates showed similar internal variability in both gene pools, with most loci being biallelic and rare alleles contributing little or marginal to overall diversity. Beta-diversity revealed that Andean and Mesoamerican groups largely share the same alleles but differ in allele frequencies, reflecting the combined effects of local selection, drift, and demographic histories.

The four subgroups identified at K=4 maintained nearly complete allelic richness, with differentiation arising mainly from differences in allele frequencies rather than unique variants. This pattern is consistent with historical farmer migration from the Andes into Amazonian areas, where likely Andean beans were introduced and later mixed with locally adapted materials, followed by continuous selection and cultivation under tropical conditions. This result is similar what was found by [55]on diversification and population structure in common beans, suggesting geographic isolation, founder effects or natural selection and introgression were involved in creating the diversity of cultivated beans.

### Population Heterozygosity

Despite the high proportion of polymorphic *loci*, observed heterozygosity (Ho ≈ 0.035–0.041) was markedly lower than expected (He ≈ 0.147–0.148). Positive and high inbreeding coefficients (FIS = 0.30–0.58) across all groups and subgroups indicate strong homozygosity, which is expected in a predominantly self-pollinating crop such as common bean. Similar patterns of heterozygote deficit have been reported in Brazil and Ethiopia [42] [28].

In both morphological and molecular characterization, no duplicates were found in the material under study, this result is similar that was found evaluating 67 accessions based on morphological traits and molecular markers from Brazil [42]; which suggests the importance of keeping all accessions in the germplasm bank since they represent a valuable source of genetic diversity. Importantly, this variability has remained largely unexploited in breeding programs, which have focused on disease resistance; lagging to obtain high yielding and high-quality cultivars in major production regions [56] [57] Incorporating Amazonian diversity into breeding pipelines could broaden the adaptive base of common bean. Hence, the conservation and systematic use of this germplasm are of strategic importance for Peru national genetic resources and future crop improvement.

## Conclusion

This study demonstrates that common bean (*Phaseolus vulgaris* L.) landraces from the Peruvian Amazon harbor remarkable morphological and genetic diversity, confirming the region as a key reservoir of agrobiodiversity. Both morphological and SNP-based analyses consistently revealed two major groups corresponding to the Andean and Mesoamerican gene pools, each subdivided into local subgroups that include unique Amazonian accessions.

Genetic diversity analyses showed similar internal variability between groups, with differences arising mainly from allele frequency rather than different allele presence, and confirmed the expected high levels of inbreeding in this predominantly self-pollinated crop. Importantly, no redundant accessions were detected, highlighting the unique value of each collected sample.

The heterozygosity analysis showed that both groups have a high proportion of polymorphic loci but very low observed heterozygosity (Ho) compared to the expected heterozygosity (He). The positive and high FIS (0.30–0.58) in all cases suggest inbreeding, which is expected on common bean.

The integration of morphological descriptors with DArTseq SNP markers proved effective in identifying structure and patterns of variation, demonstrating the complementarity of these approaches for germplasm management. Because much of this diversity is maintained only *in situ* by local farmers, there is an urgent need to reinforce both *in situ* and *ex situ* conservation strategies.

These findings underline the strategic importance of Amazonian bean landraces for national and international germplasm collections. They provide novel opportunities for breeding programs seeking traits related to tropical adaptation, resilience to climatic stresses, and farmer- and market-preferred characteristics. Harnessing this diversity is essential to ensure food security and sustainable agriculture development in the face of future challenges.

## Acknowledgements

This work was supported, in whole by Peruvian government funding-PROCIENCIA, E041 - Proyectos de Investigación Básica y Aplicada (PE501079155-2022), through the “Genómica del frijol Phaseolus spp. asociada con la diversidad cultural y ecosistémica en la Amazonia peruana” project. The authors are thankful to SAGA-Genetic Analysis Service for Agriculture, CIMMYT-Mexico; for their support for genotyping. Also, we thank local farmers for their cooperation and the research institutions (UNALM, UNSM, UNU) that supported this study.

## Author contributions

JT, MB, DV and MP contributed to conceptualization, to methodology and to writing; JP, JC and YV contributed to investigation; DV, RR and CP to data curation, writing—original draft preparation, visualization, and formal analysis; JS contributed to writing—review; JE contributed to review; RO contributed to writing—review and editing; RB contributed to conceptualization, to methodology to supervision and to funding acquisition. All authors have read and agreed to the published version of the manuscript.

## Conflicts of interest

The authors declare no conflict of interest.

## Ethical approval

This article does not contain any studies with human participants and animals and ethical approval is not applicable.

## Data availability

Supplementary Tables 1 and 2 provide the data on morphological characterization and DArTseq single nucleotide polymorphisms. They are available at https://eur03.safelinks.protection.outlook.com/?url=https%3A%2F%2Fdrive.google.com%2Fdrive%2Ffolders%2F1tqK1DZubMbASzOzGwMe3t2yRl2HDZLAz%3Fusp%3Dshare_link&data=05%7C02%7Crodomiro.ortiz%40slu.se%7Cb38dccfe9a0e4bc47d4d08dde9b01d99%7Ca3b5f0710e4947a0a40e9b7c9c4d647e%7C1%7C0%7C638923664841418090%7CUnknown%7CTWFpbGZsb3d8eyJFbXB0eU1hcGkiOnRydWUsIlYiOiIwLjAuMDAwMCIsIlAiOiJXaW4zMiIsIkFOIjoiTWFpbCIsIldUIjoyfQ%3D%3D%7C0%7C%7C%7C&sdata=jQ0ObbddyFf6LxKIbE2ik6nNmxqMXnJG3QoL%2FNLRG7E%3D&reserved=0

## Notes

### Competing Interest Statement

The authors have declared no competing interest.

